# Established rodent community delays recovery of dominant competitor following experimental disturbance

**DOI:** 10.1101/503078

**Authors:** Erica M. Christensen, Gavin L. Simpson, S.K. Morgan Ernest

## Abstract

Human activities alter processes that control local biodiversity, causing changes in the abundance and identity of species in many ecosystems. However, restoring biodiversity to a previous state is rarely as simple as reintroducing lost species or restoring processes to their pre-disturbance state. Theory suggests that established species can impede shifts in species composition via a variety of mechanisms, including direct interference (e.g. territoriality), preempting resources, or habitat alteration. Here we use a long-term experimental manipulation of a desert rodent community to examine differences in the recolonization dynamics of a dominant competitor (kangaroo rats of the genus *Dipodomys*) when patches were already occupied by an existing rodent community relative to when patches were empty. Recovery of kangaroo rat populations was slow on plots with an established community of other rodent species, taking approximately two years. In contrast, recovery of kangaroo rat populations was rapid on empty plots with no established residents (approximately 3 months). We found little evidence that the delay in kangaroo rat colonization was due to direct interference from competitors, or could be explained by differences in habitat, implicating resource preemption by the established community as the most likely mechanism. These results demonstrate that the presence of an established alternate community inhibits recolonization by new species, even those that should be dominant in the community. This has important implications for understanding how biodiversity may change in the future, and what processes may slow or prevent this change.

**Significance statement:** Ecological communities are changing due to human activities altering the processes governing local biodiversity. However restoring these processes often fails to restore the previous biodiversity state, implying that additional mechanisms contribute to community dynamics. Here we use an experimental manipulation of a desert rodent community—in which dominant competitors (kangaroo rats) were removed and then reintroduced years later—to show that the presence of previously-established species alters the dynamics of the dominant competitor’s recovery. Kangaroo rat populations took two years to recover on patches where inferior competitors were already established, compared to three months on uninhabited patches. This suggests that priority effects and initial conditions are critical to consider when predicting community response to disturbance, or in ecological restoration projects.

## Introduction

Biodiversity in many ecosystems is changing in response to anthropogenic impacts (Dornelas et al., 2014; Hillebrand et al., 2008; McGill et al., 2015), making it critical to understand the processes that accelerate change or impede our ability to reverse it. At the local scale, three main classes of interacting processes influence biodiversity: dispersal between patches (Leibold et al., 2004), environmental conditions (Chase and Leibold, 2003), and species interactions (Chesson, 2000). Alterations to any of these processes can alter local biodiversity. Dispersal links habitat patches, providing immigrants that can either rescue resident populations in danger of local extinction or introduce new species better suited to the local environment (Leibold et al., 2004). Changes in environmental conditions not only affect the physiological performance of resident species (Niehaus et al., 2012; Sears, 2005), but may also affect the outcome of competitive interactions (Davis et al., 1998) according to how well a particular habitat fulfills a species’ niche requirements (e.g., abiotic tolerances, resource requirements). The network of species interactions (e.g., competitive interactions, predation pressures, and disease susceptibilities) further restricts which species are found in which patches, even if all species can reach and survive in all the patches on a landscape. Shifts in these processes, especially shifts caused by human activities including reduced connectivity of patches (e.g., habitat fragmentation), landscape conversions (e.g., forest to pastures), and exotic species introductions are blamed for much of the biodiversity change currently being observed (Gaston et al., 2003; Haddad et al., 2015; McGeoch et al., 2010).

Theoretically, restoring biodiversity should be as simple as restoring processes to their previous, pre-disturbance state. This is the approach often employed in restoration ecology, in the form of species re-introductions (Gibbs et al., 2008) or exotic species removal (Glen et al., 2013), restoration of natural disturbance regimes to maintain suitable environments (e.g., fire regimes; Glitzenstein et al., 1995), or creating corridors to link habitat fragments (Burkart et al., 2016). After decades of attempting to restore biodiversity through these interventions it has become clear that despite our understanding of the processes controlling biodiversity, restoring biodiversity to a particular state is often more difficult than we would expect. This observation has led to the concept of alternative stable states (Beisner et al., 2003)—states or configurations of biodiversity and ecosystem processes that, once reached, are difficult to change. Altering biodiversity may be difficult once an ecosystem has settled into a new stable state due to reinforcing processes that operate to maintain the new state of the system. Thus, returning an ecosystem to a prior biodiversity state may not be as simple as just restoring major biodiversity maintenance processes to their previous levels.

Species interactions, often in combination with dispersal or environmental change, are frequently implicated in maintaining biodiversity in a particular configuration, or species composition, and impeding change (Suding et al., 2004). When a community has reached a stable state, reinforcing mechanisms work to maintain this state (Chesson, 2000) making it difficult for new species to invade an existing community (Levine et al., 2004). Inferior competitors can even prevent superior competitors from invading a community provided the inferior competitor has high initial abundance and displays interference mechanisms (e.g. territoriality in animals, overgrowth in plants) (Amarasekare, 2002). Species interactions can also impede change through indirect mechanisms. For example, species often impact their environment through their daily activities, which can help make the environment better for themselves and less suitable for other species (i.e., ecosystem engineers; Jones et al., 1997). Once removed, restoring a species later may be difficult because their absence (and perhaps the presence of other species) has altered the environment in ways that reduce that species’ ability to thrive. Whether through priority effects or habitat modification, species interactions have a strong potential to mediate the ability of biodiversity to move from one state to another and the speed at which these changes occur.

While species interactions may impede the ability of a community to return to a previous state, demonstrating whether a change in biodiversity state has actually been reversed in a natural system is complicated. Assessing state change reversal involves comparison to a “reference” state—the state that we expect our altered system to return to. Observational studies often refer to the state of the system before a specific process changed, or a spatially distinct “untouched” location as the reference state (White and Walker, 1997). However, it is often the case that more than one process is changing over time or space (e.g. climate, extinctions, patch connectivity; Rocha, 2010), making it unclear whether failure to converge to the reference state is due to the irreversible nature of the state change or simply because the reference state differs in other diversity-generating processes (Carpenter et al., 2011; Scheffer and Carpenter, 2003). To rigorously assess whether changes to an ecosystem are reversible requires replicated, unmanipulated reference plots which exhibit the expected biodiversity state under natural conditions, interdigitated with experimental plots which we have altered in some way and are trying to return to the control state (Boettiger and Hastings, 2012). These reference plots exhibit the expected state of the system given the current state of all of the processes—including the process of interest—and therefore comparisons to experimental plots provide a direct assessment of whether reversing drivers also reverses ecosystem state or yields alternative assemblages.

We examined the role of species interactions in impeding biodiversity change using a replicated multi-decadal long-term experiment that manipulated the dispersal of seed-eating rodents into experimental plots (Figure 1). The three levels of our manipulation (rodent removals, kangaroo rat removals, and controls) created a landscape with plots containing different rodent communities ranging from undisturbed (controls), to lacking a single genus of behaviorally-dominant seed eaters (kangaroo rats of genus *Dipodomys*), to consisting of only a few transient individuals (rodent removals). Many of our rodent species are territorial and sequester resources in caches, providing exactly the scenario where inferior competitors may be expected to delay colonization of a dominant species. Additionally, because kangaroo rats are important granivores in this system, their removal affects the annual plant community which serves as their resource base (Heske et al., 1993; Samson et al., 1992). In 2015, we converted half of our kangaroo rat and rodent removal plots to controls, allowing rodents to recolonize. We compared the recovery on the newly opened rodent removal and kangaroo rat removal treatments to our unchanged long-term control plots using GAMs (Generalized Additive Models) to assess whether there were differences in the dynamics of the re-invading kangaroo rats on these different types of plots. We also examined the dynamics of the other seed-eating rodents and metabolic flux of the entire rodent community to assess the roles of direct and indirect species interactions in explaining the re-colonization dynamics of kangaroo rats. Finally, we examined differences in plant species composition among the treatments to assess the impact of differences in the plant community caused by manipulating the rodent community.

**Figure 1:**
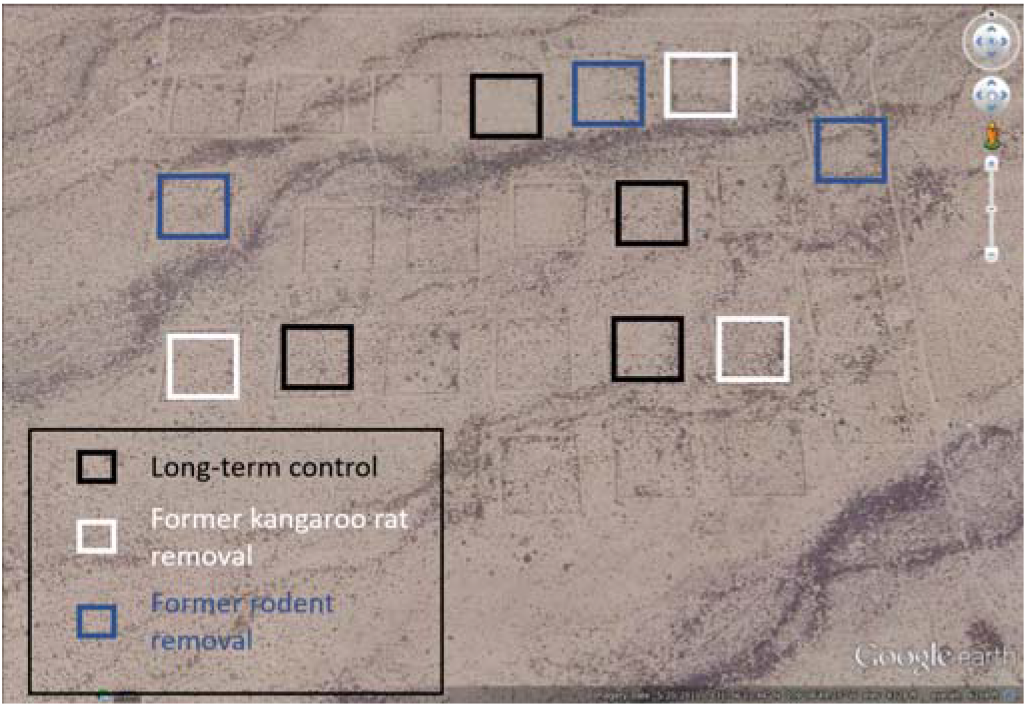
Map of the study area showing 10 experimental plots used for these analyses. The outlines of 14 additional plots not used for these analyses, but which were subjected to similar experimental manipulations, can be faintly seen. Map data from Google earth.

## Results and Discussion

### Dynamics of recovering kangaroo rat populations

Recovery of kangaroo rat populations occurred more slowly on plots containing a pre-existing rodent community than on plots where rodents were not already present (Figure 2A). We computed differences between the treatment-specific smooths from the GAM (Figure 2B), showing that kangaroo rat abundances on former rodent removals converged to control levels within 3 months (i.e., the 95% credible interval on the difference between the control and rodent removal models overlapped 0 starting June 2015). However, on the former kangaroo rat removal plots, convergence in kangaroo rat abundances did not occur until March 2017 (Figure 2B). Thus, while kangaroo rats “recovered” to control levels quickly on former rodent removals, recovery was delayed by an additional 21 months on former kangaroo rat removals. This delay does not appear to result from differences in when kangaroo rats first entered the different treatments, as kangaroo rats were detected almost immediately on both types of plots once they were opened for colonization (Figure 2A). Differences in dynamics are also unlikely to be due to differences in treatment efficacy in suppressing kangaroo rat abundances prior to being opened as controls. Both sets of treatments started with extremely low average numbers of kangaroo rats when compared to control plots (Figure 2A, also see SI Appendix: Figure S1 and Table S1). Additionally, all plots were trapped immediately prior to converting treatments to controls in 2015, ensuring that rodent removals contained no rodents and kangaroo rat removals contained no kangaroo rats at this critical point in time. Experimental characteristics also make it unlikely that differences in the recovery dynamics are due to differences in the ability of kangaroo rats to find entrances to the different treatments. All plots are embedded in the same habitat matrix where kangaroo rats are an abundant species (Figure 1), all plots have an equal number and size of gates in fences (4 gates on each side of each plot, 3.7 × 5.7 cm in size), and treatments are interspersed across the site (Figure 1). Thus, differences in the length of time required for the kangaroo rat populations to recover to control levels in the different treatments is most likely due to differences in the plots caused by the presence of a resident rodent community on the former kangaroo rat removals.

The existence of a rodent community on the kangaroo rat removal plots created a different recovery dynamic than on plots with no pre-existing established rodent community. While our results technically show that former kangaroo rat removals and controls eventually converged, it is important to note that the manner of this convergence was somewhat unusual. While former rodent removal plots converged to control plot conditions through a steady increase in kangaroo rat populations to the levels seen on control plots, former kangaroo rat removal plots converged to control conditions at a point in time when populations on control plots decreased until they reached similar levels to those on former kangaroo rat removals (Figure 2). Dynamics on the control plots were mimicked by dynamics on the former rodent removals, suggesting this decline in kangaroo rats was a general system phenomenon. However, the processes driving those declines did not similarly impact the kangaroo rats on the former kangaroo rat removals. This presents an interesting question—is the convergence between the control plots and the former kangaroo rat removal plots a real convergence where both treatments are in similar ecological states, or are the processes driving kangaroo rat abundances on these plots still different but happen to result in similar kangaroo rat populations at this moment in time? While we cannot currently answer this question, we can safely say that it took at least an additional 21 months for the former kangaroo rat removal plots to resemble control plots, though it is possible that further monitoring may lengthen this estimate.

**Figure 2:**
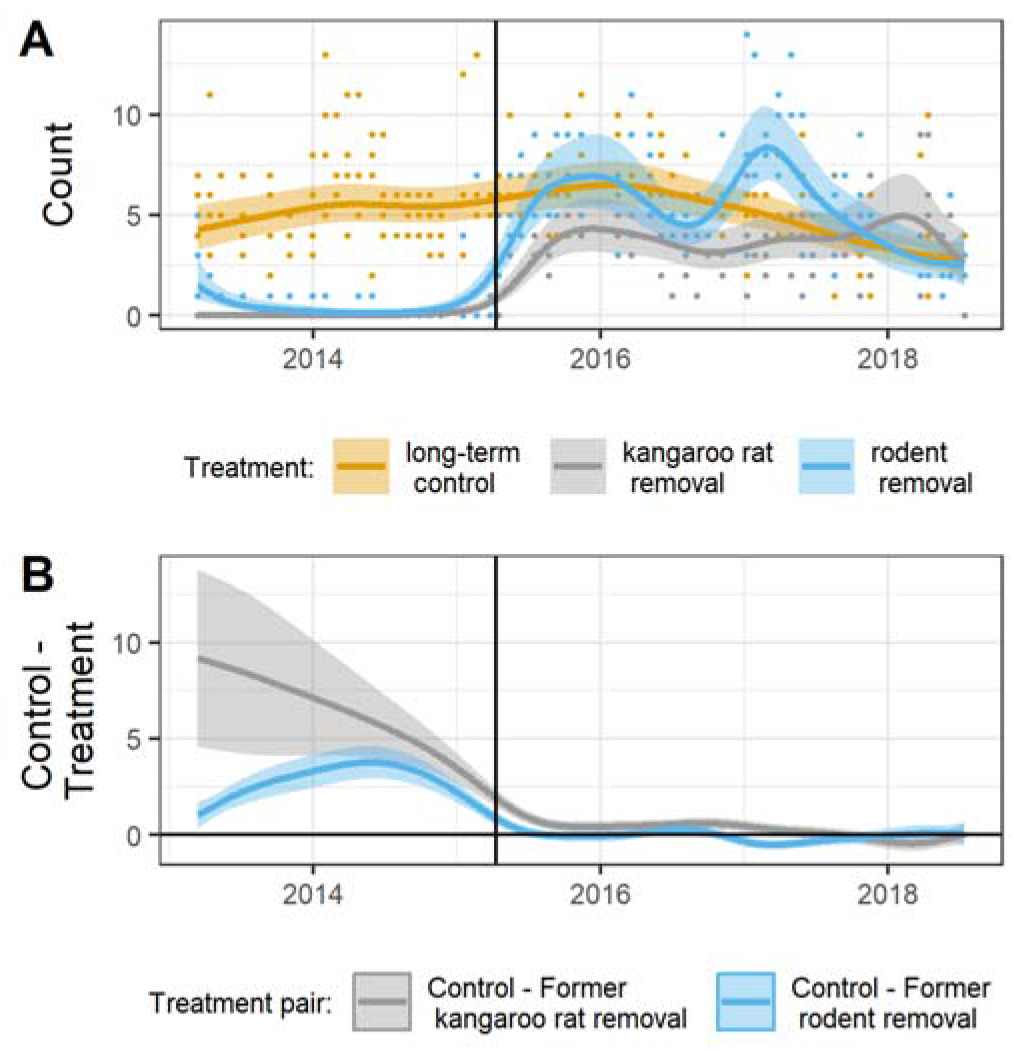
A) GAM model of number of kangaroo rats per plot and B) difference of treatment-specific smooths from GAM, shown on the link (log) scale. Vertical lines mark when treatments were changed in March 2015.

### Dynamics of non-kangaroo rat populations

If the delayed recovery of kangaroo rats was caused by direct interactions with the resident species on the former kangaroo rat removal plots, we would expect to see a slow decline in these species as resident individuals were gradually displaced from their territories by invading kangaroo rats. Instead, abundances of the non-kangaroo rat seed-eaters on both treatment types converged quickly to levels observed on controls. Before the change in treatments in 2015, abundances of the non-kangaroo rat rodents on kangaroo rat removals were higher than on controls plots (kangaroo rat removals averaged 7.8 individuals/plot/month; controls 5.7 individuals) and much lower than control levels on rodent removals (3.1 individuals/plot/month) (See SI Appendix: Figure S2 and Table S2). After all plots were converted to controls, the number of non-kangaroo rat rodents on both the former rodent removals and the former kangaroo rat removals quickly converged to control plot levels within a few months (Figure 3B). If we focus on Bailey’s pocket mouse (*Chaetodipus baileyi*), the species most likely to compete directly with kangaroo rats (Ernest and Brown, 2001; Thibault et al., 2010), we see that the abundance of this species dropped quickly on former kangaroo rat removals once kangaroo rats were reintroduced (Figure 4). In fact, Bailey’s pocket mouse became extremely rare on all plot types within 9 months of the treatment change. The rapid decline in non-kangaroo rat species is consistent with previous research showing that kangaroo rats are behaviorally dominant over the other, typically smaller, seed-eating rodents (Leaver and Daly, 2001; Reichman and Price, 1993). Because differences between treatments in non-kangaroo rat species abundances disappeared quickly post-intervention, it seems unlikely that direct interference by the non-kangaroo rat species explains the 21-month delay in the recovery of kangaroo rats on kangaroo rat removals. There is an unexplained delay of at least a year between when these smaller granivores were displaced from their territories and when kangaroo rat abundances recovered to control levels.

**Figure 3:**
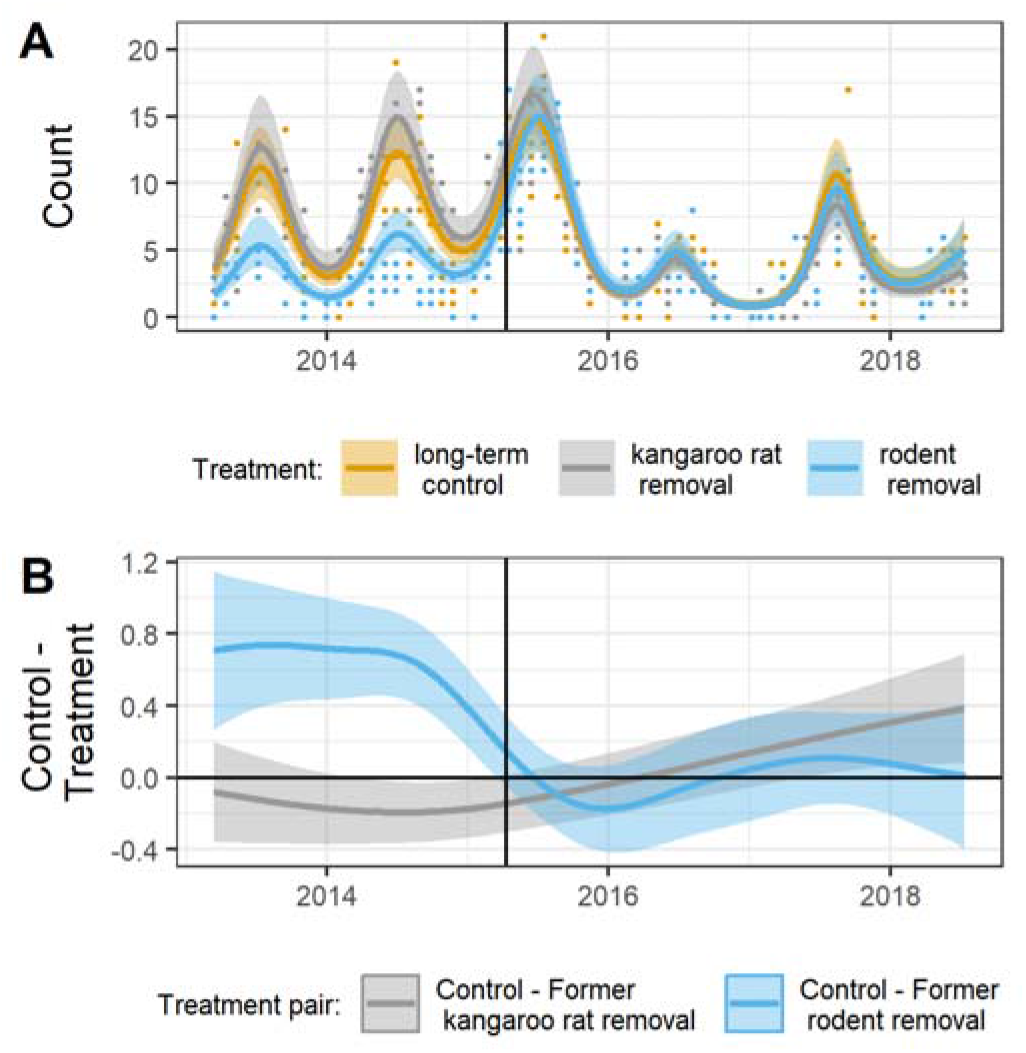
A) GAM model of abundances of non-kangaroo rat species per plot and B) difference of treatment-specific smooths from GAM, shown on the link (log) scale. Vertical lines mark when treatments were changed in March 2015.

**Figure 4:**
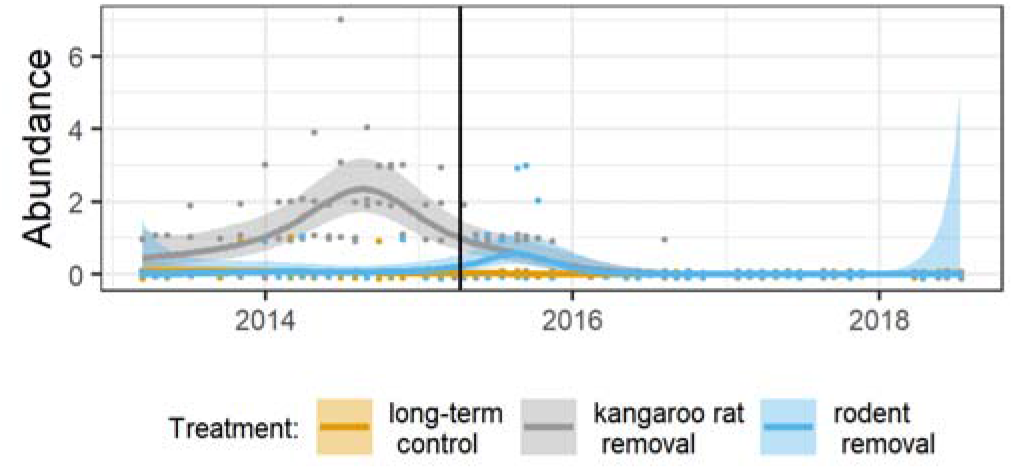
Populations of Bailey’s pocket mouse on experimental plots. Solid lines are average by treatment and month. Vertical line marks when treatments were changed in March 2015.

### Plant community differences by treatment

Differences between treatments in the ability of kangaroo rats to invade could be caused by differences in the plant communities related to the pre-2015 rodent manipulations. While rodents can impact the plant community as consumers, the plant community can also impact the rodent community through two mechanisms: via the habitat structure (Rosenzweig and Winakur, 1969) and resource base that the vegetation provides (Reichman, 1975). If the plant community impeded colonization by kangaroo rats on former kangaroo rat removals, we would expect to see strong pre-2015 differences in plant composition between the kangaroo rat removal plots and rodent removal plots, where colonization was not impeded. We conducted a pCCA by treatment for the 7 years leading up to the treatment change in 2015 (Figure 5; see Methods) to assess differences in the plant community while controlling for between-year variation. The effect of treatment was significant for both the winter plant community (pCCA permutation test: R^2^_CCA_ = 0.03 and p = 0.002) and the summer plant community (pCCA permutation test: R^2^_CCA_ = 0.04 and p = 0.004). However, in both cases the proportion of variance explained by treatment was small (less than 5%). Thus, while the experimental manipulations did impact plant composition, these impacts were only weakly different between the rodent removals and kangaroo rat removals during this time period (See Figure 5, and SI Appendix Fig. S4-S5).

**Figure 5:**
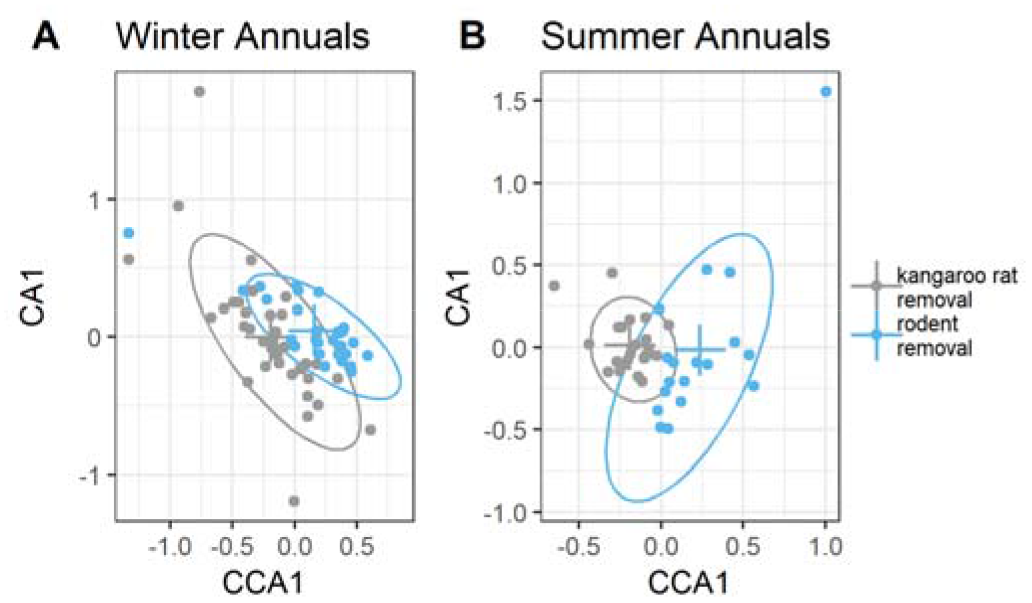
pCCA of treatment effects on plant composition for A) winter and B) summer annual plants. Crosses indicate data centroids by treatment, and ellipses enclose 95% of the data points.

### Dynamics of total metabolic flux of the rodent community

The third option for why kangaroo rats were able to invade former rodent removals more easily than former kangaroo rat removals is the possibility that resources—in this case seeds—on rodent removals were more readily available to new colonizers. Seeds may have been more plentiful on rodent removals due to lack of consumption by an established rodent community. Alternatively, while there may not have been more seeds on rodent removals, they may have been easier to find because they had not been gathered and deposited into caches (Leaver and Daly, 2001; Swartz et al., 2010). While we do not have information about the distribution or availability of seeds, we can gain some insights via examination of metabolic energy flux of the rodent community on the different treatments. Community metabolic flux (Brown et al., 2004; Padfield et al., 2018) is a measure of the energy intake rate required to sustain the rodent community on a plot, and thus is an index of resource use by the community as a whole (see Methods for calculation). If there were more resources available on rodent removals due to seed accumulation, then we would expect the metabolic energy flux on rodent removals to be higher than on controls. However, there is little evidence that the community metabolic flux was higher on the rodent removal plots than on the controls (Figure 6) except for a brief period of time beginning in 2017, which was over a year after the treatments were changed. In contrast, metabolic flux on the former kangaroo rat removals remained below both the former rodent removals and controls until 2018. The decline in the non-kangaroo rat rodents early in the recovery process (Figure 3) should have freed up resources for the kangaroo rats to use, bringing energy use on the control plots and former kangaroo rat removals closer together, as happened on the former rodent removals. However, our results suggest that there were still unused resources available on the former kangaroo rat removals that the kangaroo rats were not accessing. The fate of these missing resources is unknown. It is possible that these resources were hidden in caches that kangaroo rats were simply unable to find or that those resources were pre-empted by one of the other seed-eating taxa in this ecosystem (i.e. ants or birds). Unfortunately, we are currently unable to test either of these hypotheses.

**Figure 6:**
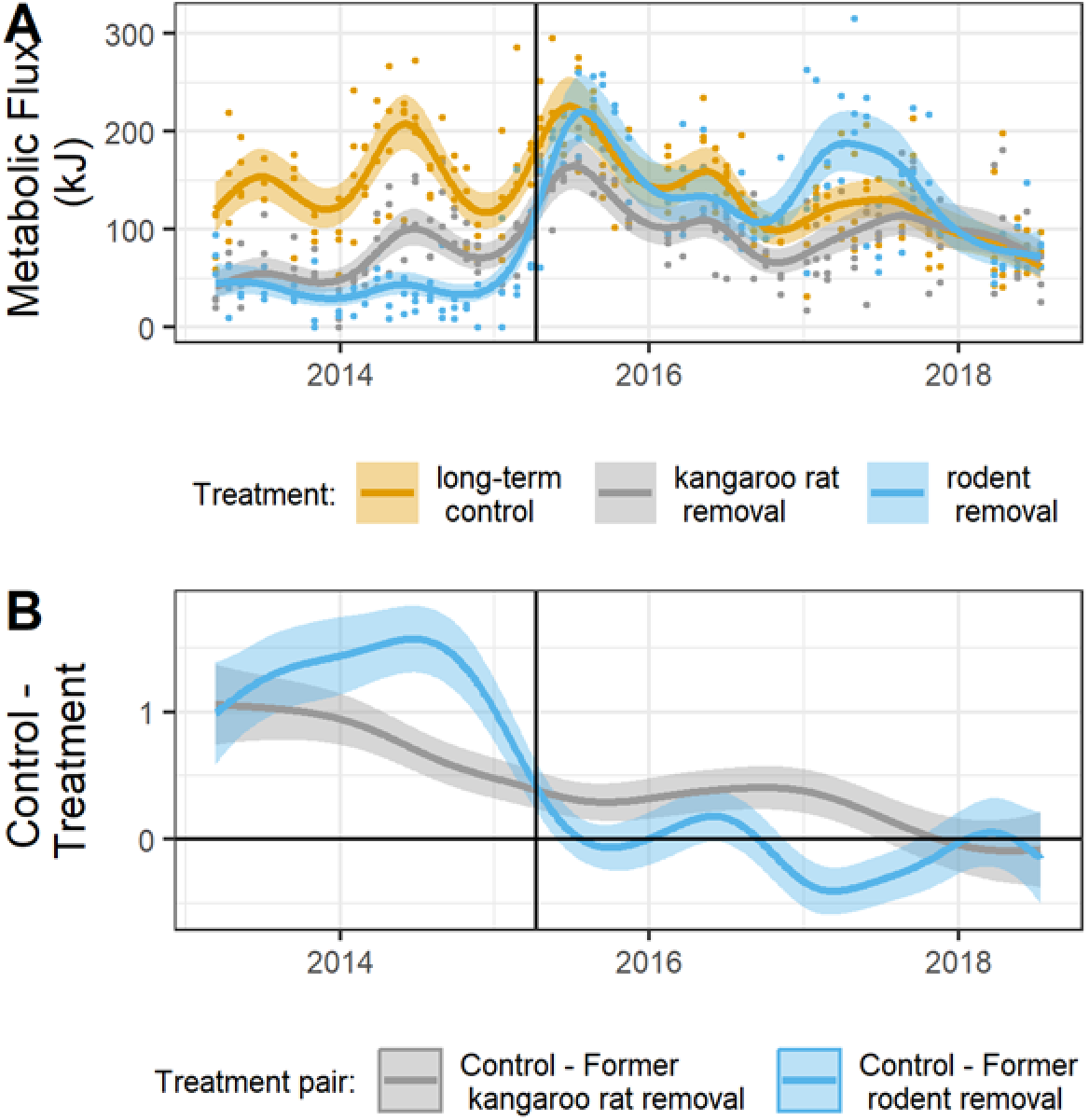
A) GAM model of community-level metabolic flux per plot and B) difference between treatment-specific smooths from GAM, shown on the link (log) scale. Vertical lines mark when treatments were changed in March 2015.

## Conclusions

The scale and nature of our experiment suggest that the ability of a resident community to impede invasion by a dominant competitor can be extremely strong. The large source pool of immigrants from the habitat matrix should have provided a strong invasion advantage to the kangaroo rats, and the relatively small plot size (50m × 50m) ensured that only dispersal (not population growth) was required for kangaroo rats to establish control-level populations on experimental plots. That this advantage still resulted in a long delay in plot convergence suggests that we should expect long transient dynamics when we attempt to shift the biodiversity state of an ecosystem (sensu Hastings, 2004). While transient dynamics may eventually lead to some expected stable state, transient dynamics can also lead to unexpected and complex dynamics, especially in the presence of stochasticity, a common feature of ecological systems (Hastings, 2004; Hastings et al., 2018).

Assessing the reversibility of state change is difficult because ecosystems are dynamic, meaning that the “expected” state of the system may vary over time. Without a concurrent reference state to compare to, the inherent dynamics of an ecosystem can create the appearance of convergence (i.e. treatments become similar to a control state from the past, but are not convergent with the control state of the present). Even with concurrent reference plots, assessing whether two ecosystem states are similar is potentially tricky, as shown with our unexpected convergence driven by control plots becoming more similar to former kangaroo rat removals instead of vice versa. While having concurrent reference plots is the ideal design, implementation of replicated experiments at a scale where alternative states can be generated is challenging, which explains why few studies of state changes in ecology (e.g., regime shifts) involve replicated experiments and fewer still occur under field conditions (Schröder et al., 2005). Even more challenging is that manipulations need to switch between ecologically relevant alternative states, which requires not only an understanding of which ecological drivers lead to alternative states, but the ability to manipulate these drivers in a controlled way. While few field systems fit all of these criteria, those that do are critical for rigorously assessing theoretical and conceptual aspects of alternative stable states and related ideas (Carpenter, 2003).

It is increasingly clear that established communities of species impact how the biodiversity of an ecosystem changes over time (Amarasekare, 2002; Fukami, 2015; Thibault and Brown, 2008). Species interactions can alter the dynamics of biodiversity recovery and impede the ability of species to return to expected population sizes when biodiversity maintenance processes, such as dispersal in our case, are restored. Even if established species are competitively inferior to invading species, the existence of an established community will have impacts on the environment and the availability and distribution of resources (Amarasekare, 2002; Grman and Suding, 2010). When biodiversity maintenance processes change, the impacts of already-established species can affect how biodiversity changes and the time it takes for these changes to occur. Our results suggest that attempts to restore biodiversity by altering a biodiversity process (i.e. dispersal in our experiment) may result in slow or unexpected dynamics due to the complicating effects of species already present in the ecosystem. Our results suggest not only that existing established species can alter or delay changes in biodiversity state, but also the reverse—that disruptions of an existing community may facilitate and/or speed up biodiversity shifts. The rapid conversion of rodent removals to controls was made possible by the absence of the influence of an existing rodent community. This result is consistent with other work that has noted rapid shifts in biodiversity states after disturbances (Christensen et al., 2018b; Smith, 2011; Turner, 2010). Thus disturbances that eliminate the impacts or advantages of an existing community may create conditions that make biodiversity state shifts more probable. Understanding the importance and prevalence of the mechanisms by which existing assemblages of species prevent or alter trajectories of change will be critical for our ability to predict both how biodiversity might change and the conditions that increase the probability of rapid shifts in biodiversity.

## Site and Methods

The experiment was conducted at the Portal Project, a 20-hectare study site, 6 km northeast of the town Portal, Arizona in the Chihuahuan desert (Ernest et al., 2018). Twenty-four 50 m by 50 m experimental plots are enclosed by a 50 cm high fence, and rodent access is controlled by gates in the fences at ground level. The size and/or absence of gates regulates access to the plot: large gates for controls, smaller gates for kangaroo rat removals, and no gates for rodent removals. Each plot is trapped every month using 49 Sherman traps, and information from captured rodents is recorded including species, sex, and weight. Any rodents caught on total removal plots, and kangaroo rats caught on kangaroo rat removal plots, are recorded and relocated at least a quarter mile away from the site. Because there are two relatively distinct periods of annual plant growth with little overlap in species, plant abundances are sampled on each plot twice each year—once to capture the summer community and again to capture the winter community. On each plot, all plants are identified and counted on sixteen 0.25 m^2^ quadrats placed at permanently marked locations.

While data collection began at this site in 1977, this analysis focuses on the time period 2013–2018. In March 2015, we changed the treatments on 12 of the 24 plots. The pre-existing treatments on those 12 altered plots had been in place continuously since 1989. We focus here on plots that were formerly kangaroo rat removal plots that had gates enlarged to become controls (3 plots), plots that were formerly total rodent removal plots that had new gates cut to become controls (3 plots). Plots that were control plots for the entire period 2013–2018 (4 plots) serve as our reference plots for assessing the dynamics of our newly-created control plots. All data and code for these analyses are available on GitHub (https://github.com/emchristensen/PlotSwitch) and archived on Zenodo (Christensen et al. 2018a).

### Time series analysis using GAMs

We used GAMs to assess the effect of treatment on various rodent community metrics over time, using the R package mgcv version 1.8–23 (Wood, 2011) for R 3.5.1 (R Core Team, 2018). We constructed a GAM for each metric including a smoothing factor based on pre-2015 treatment type, then computed the difference between pairs of treatment-specific smooths, focusing on comparing former kangaroo rat removals to long-term controls, and former rodent removals to long-term controls. Where the Bayesian credible intervals for the difference in smooths included zero, we interpreted this to mean there was no effective difference in the metric of interest for that pair of smooths.

To assess the ability of competitively-dominant kangaroo rats to colonize suitable patches, we constructed time series of pooled number of individuals of the three species of kangaroo rats found at the site (*Dipodomys merriami, D. ordii*, and *D. spectabilis*) on each plot over time. Since we expect inferior competitors to be displaced by the invasion of kangaroo rats, we also constructed time series of pooled number of individuals of non-kangaroo rat seed-eating species (10 species from 5 genera: *Baiomys taylori, Chaetodipus baileyi, C. penicillatus, Perognathus flavus, Peromyscus eremicus, P. leucopus, P. maniculatus, Reithrodontomys fulvescens, R. megalotis*, and *R. montanus*). Because these two time series consist of count data, we used a Poisson distribution in both GAMs. We found that there were some small but significant differences between plots within treatments, and so included plot-specific smooths in the GAMs as well.

To examine effects at the community level, we analyzed time series of community metabolic flux of the seed-eating rodent community on each plot. Total metabolic flux represents an estimate of community size and resource uptake of the community as a whole (Padfield et al., 2018), and is generally less variable through time than species composition or species abundances (White et al., 2004). Metabolic rate of individual organisms scales with body size according to the equation: *E* ∝ *m*^3/4^ where *E* is metabolic rate (or energy) and *m* is mass of the individual (Brown et al., 2004). We estimated metabolic flux of individual captured rodents based on measured masses, and summed by plot and time step to obtain total community metabolic flux. We fitted a GAM to the resulting time series, using a Tweedie distribution, and plot-specific smooths as well.

### Plant community analysis

Differences in plant community composition were assessed using a partial canonical correspondence analysis (pCCA) and permutational significance tests, controlling for between-year effects (using R package vegan: Oksanen et al. 2018). We square root transformed the plant abundance data to account for large differences in total abundance between years and species. Due to project funding gaps plant data was not collected in all years leading up to the treatment change in 2015. We used all data available going back to 2008, which amounted to three years of data for the summer annual community (2008, 2011, and 2014) and five years of data for the winter annual community (2008, 2012, 2013, 2014, and 2015).

## Acknowledgements

Field data collection was supported by grant DEB-1622425 from the National Science Foundation, which also supported E. Christensen. We would like to thank D. Valle, T. Palmer, and B. Baiser for their helpful comments on the manuscript. E. Christensen was also supported by funds to the USDA-ARS Jornada Experimental Range 3050–11210-009–00-D.

