## Appendix for "Established rodent community delays recovery of dominant competitor following experimental disturbance"

Supplementary Information Text

Comparison of treatments pre-2015

To better interpret the effect of the treatment change, we compared the three metrics of interest (number of kangaroo rat individuals, number of non-kangaroo rat individuals, and total metabolic flux) between treatments for the 2-year time period leading up to the treatment change. We used Generalized Linear Mixed Models (GLMMs) to analyze the effect of treatment on each of the three metrics while accounting for between-plot differences. Models of numbers of individuals were fit with Poisson distributions, and the model of total metabolic flux was fit with a Tweedie distribution. All analyses were conducted using mgcv in R (version 3.5.1; R Core Team 2018) with post hoc tests using the glht() function in the multcomp package (version 1.4-8; Hothorn et al. 2016). There were significant pair-wise differences between all pairs of treatments for number of kangaroo rat individuals, number of non-kangaroo rat individuals, and total metabolic flux (Tables S1-S3). Thus we expect the experimental treatment changes in 2015 to have measurable effects on these metrics.

Additional plant community comparisons

If the slight differences in plant composition made kangaroo rat removals more different from controls than the rodent removals were, then perhaps these differences could contribute to the delayed response on former kangaroo rat removals as kangaroo rats dealt with an environment that was slightly more alien than the one they were accustomed too. However, there is no evidence that this was the case. In fact the evidence suggests the opposite—when comparing rodent removals to controls, the treatment effect explains a slightly higher fraction of the variation in plant species composition (pCCA permutation test: winter annuals, R^2^_CCA_ = 0.06 and p = 0.002; summer annuals, R^2^_CCA_ = 0.05 and p = 0.002) than it does for the comparison between kangaroo rat removals and controls (pCCA permutation test: winter annuals, R^2^_CCA_ = 0.02 and p = 0.002; summer annuals, R^2^_CCA_ = 0.03 and p = 0.004; Figure S4-S5). This suggests that the plant community on the rodent removals was slightly more different from the control plots. While this does not rule out the possibility that the plant community negatively impacted the ability of kangaroo rats to reinvade former kangaroo rat removal plots, the support for this as a mechanism is weak.

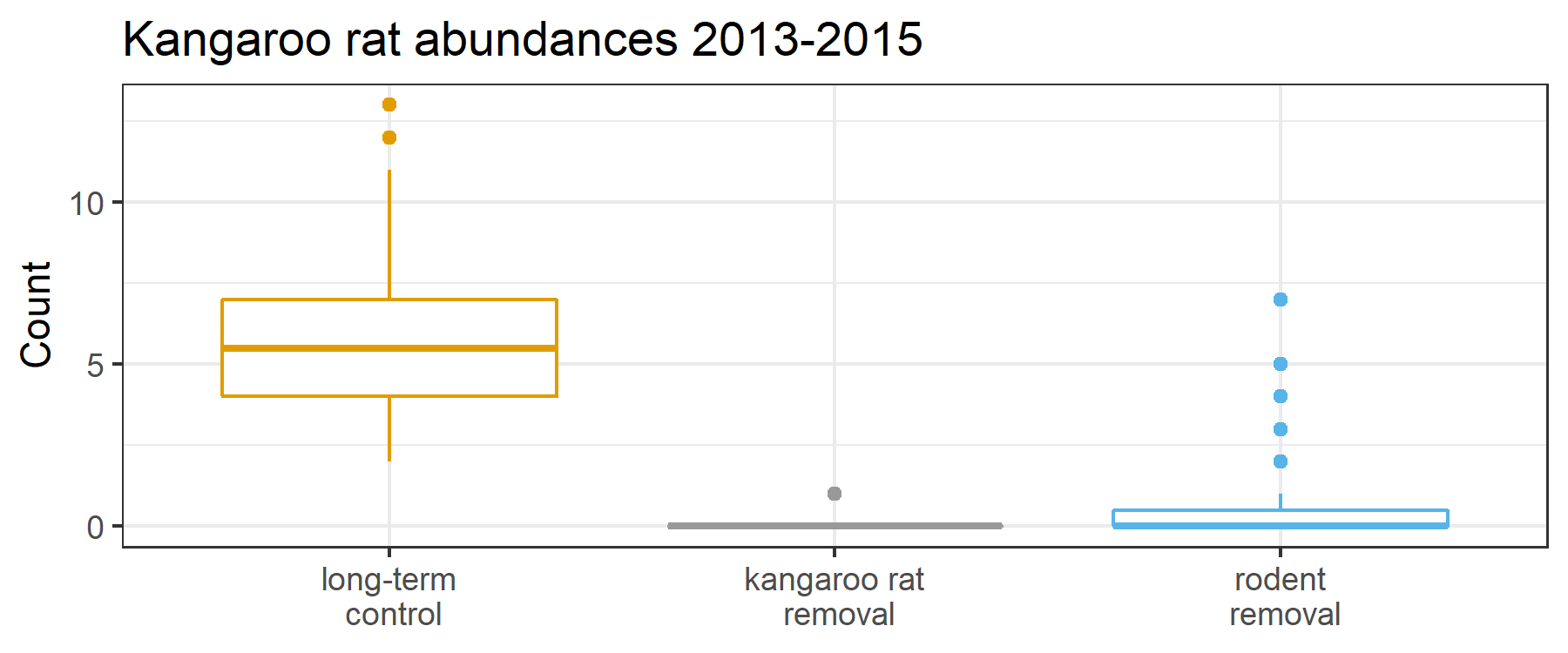

Fig. S1. Number of kangaroo rat individuals captured on plots by experimental treatment for the period before the treatment change, 2013-2015.

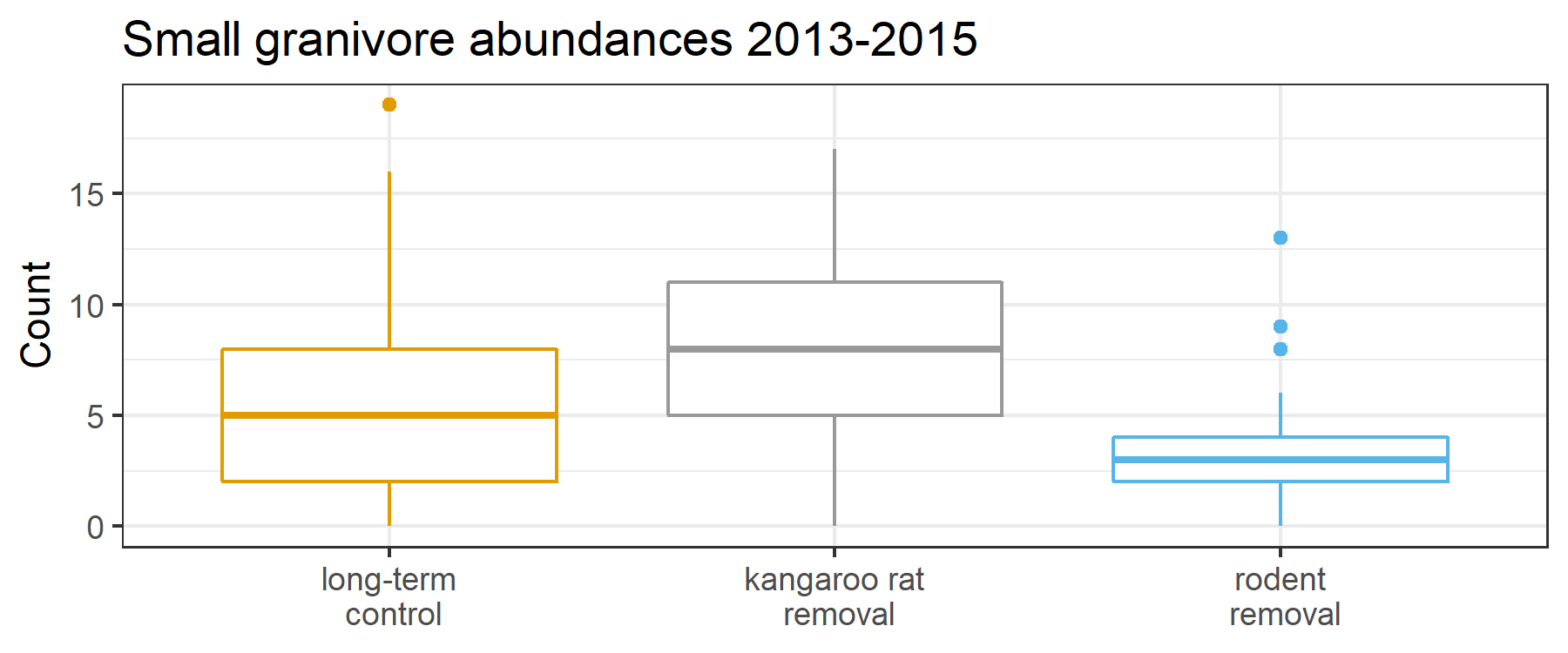

Fig. S2. Number of individuals of non-kangaroo rat species captured on plots by experimental treatment for the period before the treatment change, 2013-2015.

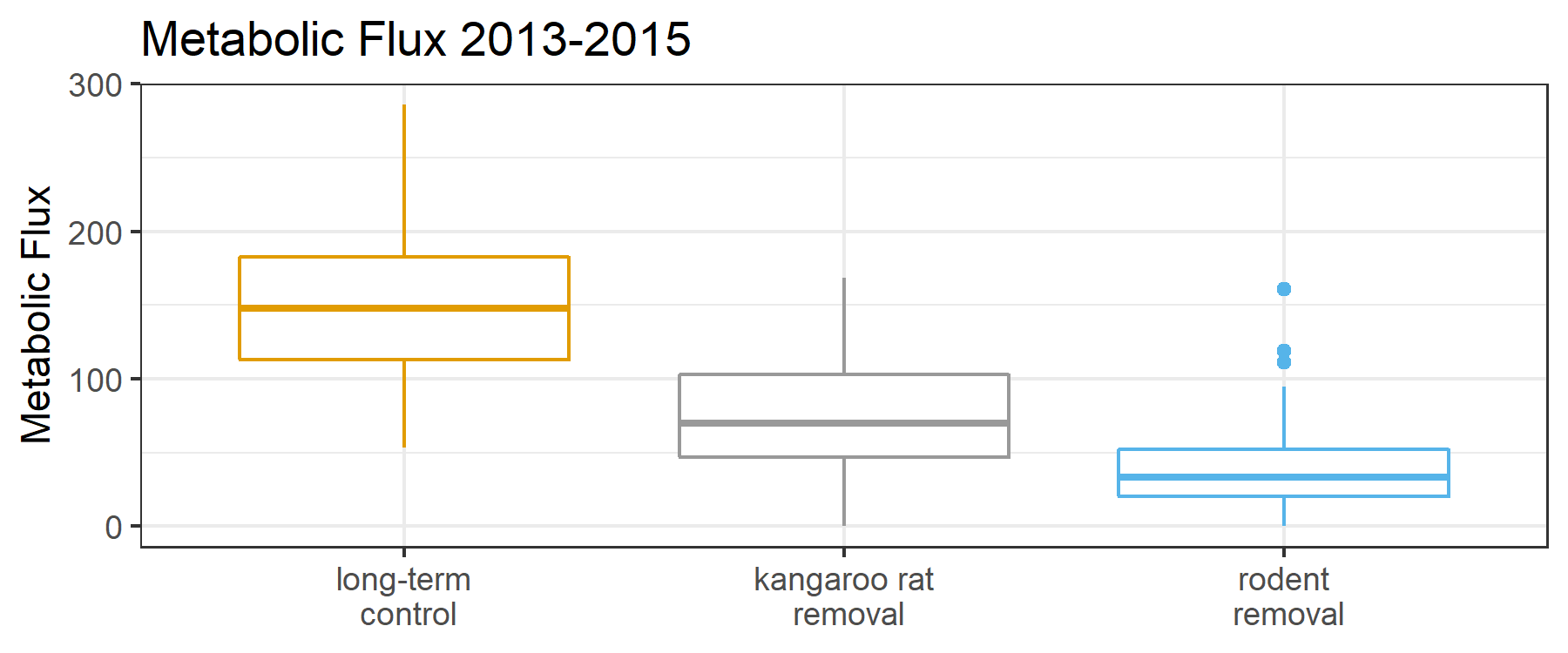

Fig. S3. Total metabolic flux of the entire rodent community captured on plots by experimental treatment for the period before the treatment change, 2013-2015.

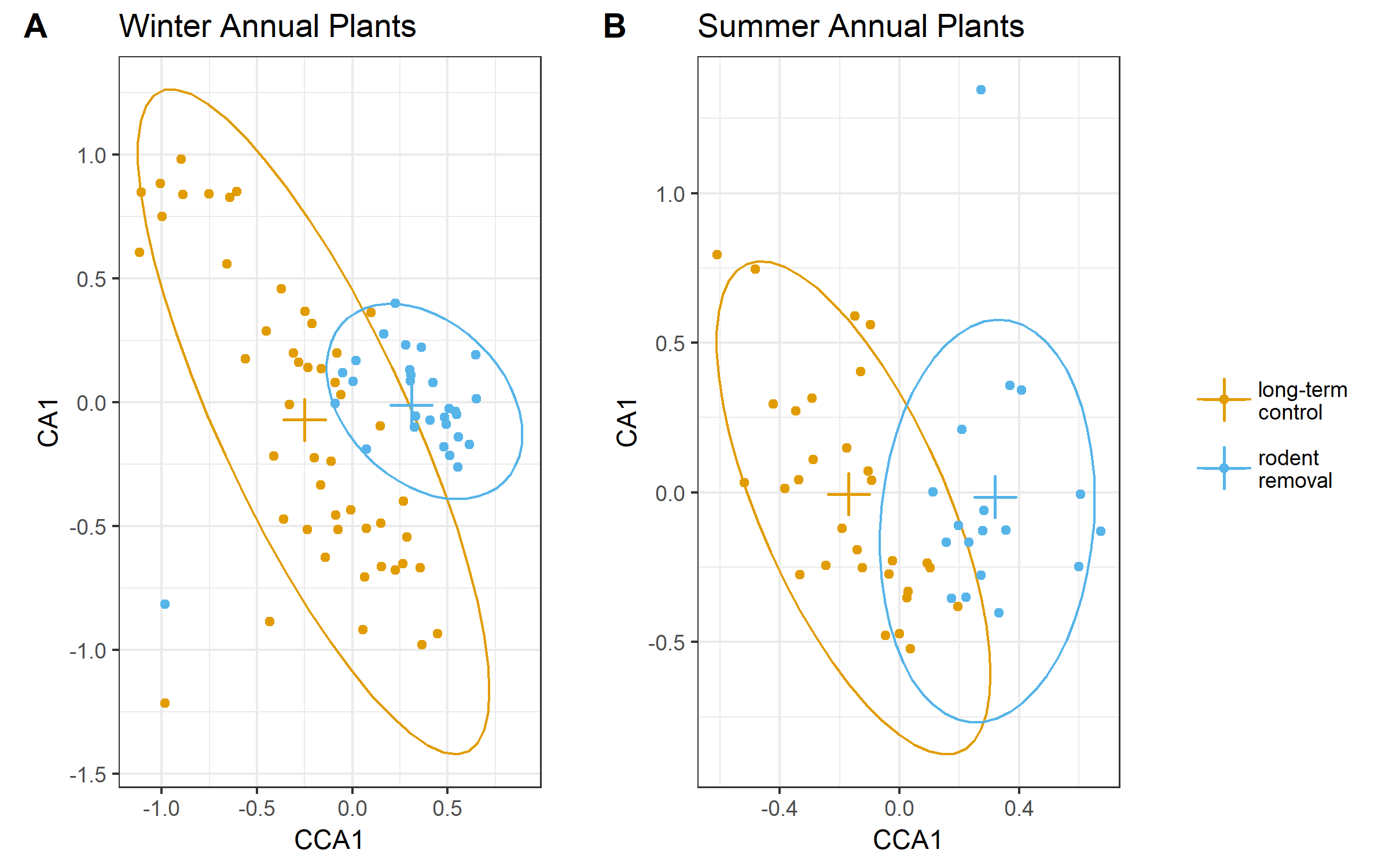

Fig. S4. Comparison of plant communities (summer and winter) on controls to plant communities on rodent removals before the treatment change in 2015.

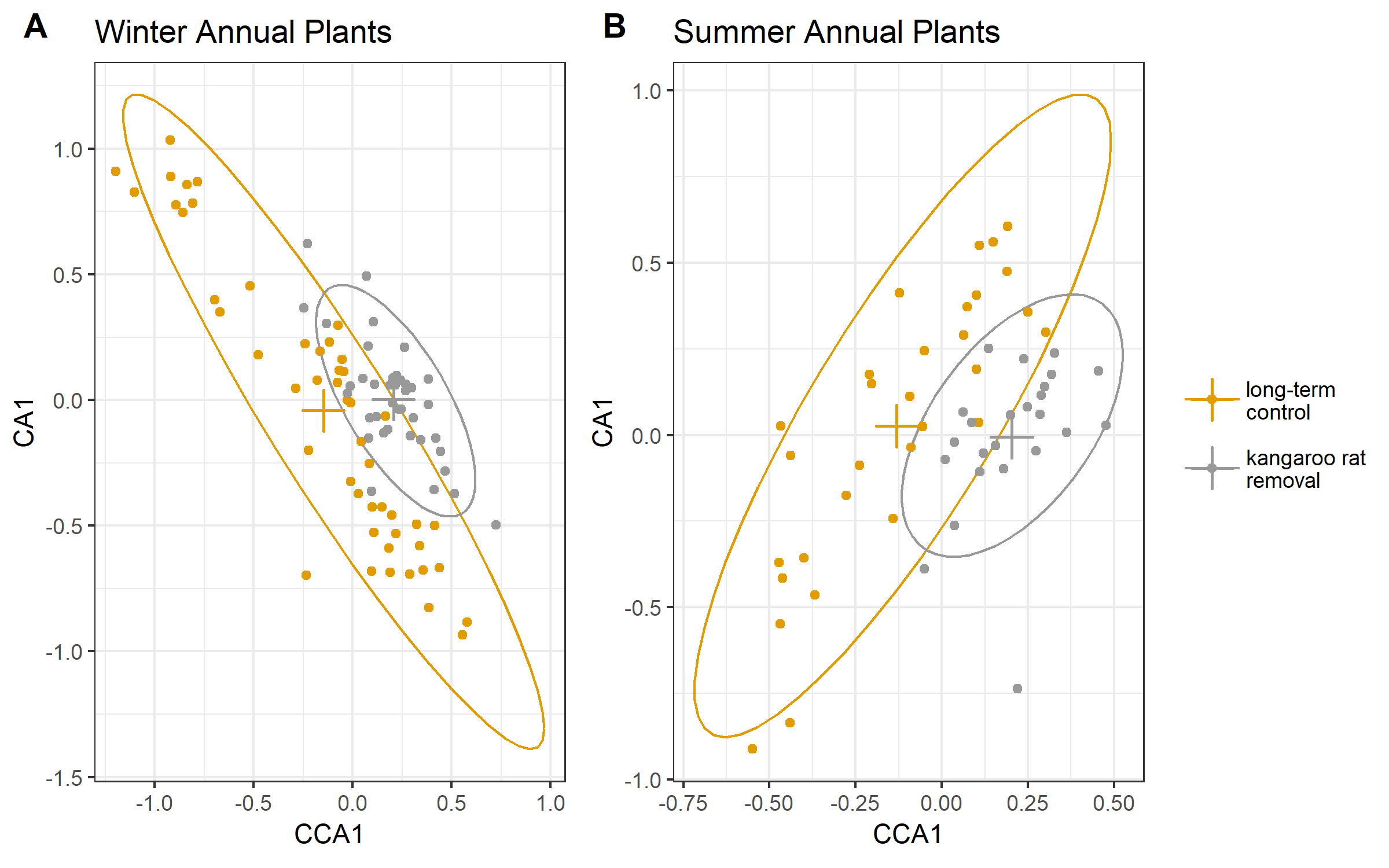

Fig. S5. Comparison of plant communities (summer and winter) on controls to plant communities on kangaroo rat removals before the treatment change in 2015.

Table S1. Multiple comparisons of means: Tukey contrasts for number of kangaroo rat individuals 2013-2015.

| Treatment pair | Estimate | Std. error | Z | P |
| --- | --- | --- | --- | --- |
| Kangaroo rat removal – control | -5.906 | 1.003 | -5.890 | < 0.001 |
| Rodent removal – control | -2.441 | 0.192 | -12.713 | < 0.001 |
| Rodent removal – kangaroo rat removal | 3.465 | 1.018 | 3.406 | 0.0015 |

Table S2. Multiple comparisons of means: Tukey contrasts for number of non-kangaroo rat individuals 2013-2015.

| Treatment pair | Estimate | Std. error | Z | P |
| --- | --- | --- | --- | --- |
| Kangaroo rat removal – control | 0.3210 | 0.1346 | 2.384 | 0.0449 |
| Rodent removal – control | -0.5888 | 0.1455 | -4.047 | < 0.001 |
| Rodent removal – kangaroo rat removal | -0.9098 | 0.1519 | -5.989 | < 0.001 |

Table S3. Multiple comparisons of means: Tukey contrasts for total metabolic flux 2013-2015.

| Treatment pair | Estimate | Std. error | Z | P |
| --- | --- | --- | --- | --- |
| Kangaroo rat removal – control | -0.70047 | 0.07192 | -9.740 | << 0.001 |
| Rodent removal – control | -1.33635 | 0.08886 | -15.039 | << 0.001 |
| Rodent removal – kangaroo rat removal | -0.63588 | 0.10037 | -6.335 | << 0.001 |
